# Microtubule glycylation promotes basal body attachment to the cell cortex

**DOI:** 10.1101/620476

**Authors:** Anthony D. Junker, Adam W. J. Soh, Eileen T. O’Toole, Janet B. Meehl, Mayukh Guha, Mark Winey, Jerry E. Honts, Jacek Gaertig, Chad G. Pearson

**Affiliations:** Department of Cell and Developmental Biology, University of Colorado School of Medicine, Aurora, CO 80045.; Department of Molecular, Cellular and Developmental Biology, University of Colorado, Boulder, CO 80302; Department of Molecular and Cellular Biology, University of California, Davis, CA 95616.; Department of Biology, Drake University, 2507 University Avenue, Des Moines, IA 50311.; Department of Cellular Biology, University of Georgia, Athens, GA 30602

**Keywords:** Cilia, basal bodies, microtubule, ciliary array, multiciliated, tubulin post-translational modifications.

## Abstract

Motile cilia generate directed hydrodynamic flow that is important for the motility of cells and extracellular fluids. To optimize directed hydrodynamic flow, motile cilia are organized and oriented into a polarized array. Basal bodies (BB) nucleate and position motile cilia at the cell cortex. Cytoplasmic BB-associated microtubules are conserved structures that extend from BBs. Using the ciliate, *Tetrahymena thermophila*, combined with EM-tomography and light microscopy, we show that BB-appendage microtubules assemble coincident with new BB assembly and are attached to the cell cortex. These BB-appendage microtubules are specifically marked with post translational modifications of tubulin, including glycylation. Mutations that prevent glycylation shorten BB-appendage microtubules and disrupt BB positioning and cortical attachment. Consistent with the attachment of BB-appendage microtubules to the cell cortex for BB positioning, mutations that disrupt the cellular cortical cytoskeleton similarly disrupt the cortical attachment and positioning of BBs. In summary, BB-appendage microtubules promote the organization of ciliary arrays through attachment to the cell cortex.

**SUMMARY STATEMENT:** Basal bodies position motile cilia at the cell cortex. This study finds tubulin glycylation to promote BB-associated microtubule elongation and structural attachment of basal bodies to the cell’s cortical cytoskeleton.

## INTRODUCTION

Motile cilia produce directed fluid flow for fundamental biological activities, including cell movement through aqueous environments and mucociliary clearance in respiratory airways. To maximize fluid flow, cells employ hundreds of motile cilia arranged into polarized arrays. Each cilium beats in a coordinated and asymmetric pattern to generate fluid flow. Cilia are nucleated, positioned and oriented at the cell cortex by basal bodies (BB). BBs comprise a cylinder of nine symmetrically arranged triplet microtubules. The distal ends of BBs attach to the cell cortex through fibers that symmetrically emanate from the ends of each triplet microtubule (Burke et al., 2014, Siller et al., 2017, Yang et al., 2018). Additional BB-associated structures project asymmetrically from BBs, linking them to other cytoskeletal elements within the cell cortex (Herawati et al., 2016, Kunimoto et al., 2012, Tateishi et al., 2017, Turk et al., 2015, Iftode et al., 1996). Such linkages ensure BB organization and orientation even in the face of forces produced by motile cilia (Galati et al., 2014, Werner et al., 2011, Mahuzier et al., 2018). Thus, cilia position and polarity are directed by the linear organization and orientation of BBs.

Microtubules associated with BBs promote cilia organization. In mouse ciliated epithelial cells, BB-associated microtubules create a network of interconnected BBs that organize and maintain the proper spacing and orientation of cilia (Herawati et al., 2016, Kunimoto et al., 2012). BB-associated microtubules in *Xenopus* multi-ciliated epithelia of skin ensure uniform spacing between BBs, which is important for effective hydrodynamic flow (Werner et al., 2011, Elgeti and Gompper, 2013). How BB-appendage microtubules connect to the rest of the cell is not yet known.

The tubulin subunits in BBs, cilia and BB-associated microtubules have post-translational modifications (PTMs), including acetylation, glutamylation and glycylation (Gadadhar et al., 2017a). Microtubule PTMs contribute to microtubule plus-end dynamics, force production by motor proteins and mechanical properties of microtubules (Gadadhar et al., 2017a, Janke, 2014). Acetylation occurs by the enzymatic activity of the MEC17 / α-TAT acetyltransferase at the K40 residue of α-tubulin located in the lumen of microtubules (Akella et al., 2010, Shida et al., 2010). Most other tubulin modifications occur at α- and β-tubulin C-terminal tails that are exposed on the microtubule outer surface. Polymeric tail modifications are catalyzed by the TTLL (Tubulin Tyrosine Ligase-Like) family of enzymes (Wloga et al., 2009, Janke et al., 2005, Rogowski et al., 2009). Microtubule glycylation is initiated by the addition of a glycine to a glutamic acid side chain in the tubulin C-terminal tail sequence (Redeker et al., 1994). This monoglycylation is catalyzed by Ttll3 or Ttll8. Monoglycylation may be followed by the addition of multiple glycine residues, which is catalyzed by the polyglycylase, Ttll10 (Rogowski et al., 2009, Wloga et al., 2009). Polyglycylases require monoglycylated microtubules for their substrate; when monoglycylation is absent, microtubules are not glycylated (Ikegami and Setou, 2009, Rogowski et al., 2009). Ttll3-dependent glycylation promotes cilia formation and elongation presumably by affecting properties of axonemal microtubules (Bosch Grau et al., 2017, Wloga et al., 2009, Gadadhar et al., 2017b). Morpholinos targeting zebrafish Ttll3 disorient cilia in pronephric ciliary arrays suggesting that tubulin glycylation is important for either BBs or cilia (Pathak et al., 2011). How glycylation promotes cilia orientation is unknown.

Like vertebrates and other protists, *Tetrahymena* cells have BB-associated microtubules or BB-appendages that are important for organizing and orienting cilia and BBs into polarized arrays. BB-appendage microtubules are post-translationally modified, however, a role for these modifications in organizing or orienting cilia and BBs has not been identified (Callen et al., 1994, Akella et al., 2010, Tassin et al., 2015, Wloga et al., 2008). BBs are endowed with three BB-appendage structures: striated fibers (SF), transverse microtubules (tMT), and post-ciliary microtubules (pcMT). SFs extend towards the cell anterior and establish and maintain BB organization and orientation (Galati et al., 2014, Jerka-Dziadosz et al., 1995). tMTs and pcMTs are composed of microtubule bundles that nucleate from the BB base and extend transversely and posteriorly, respectively, toward the cell’s cortical cytoskeleton (Fig. 1). The consistent geometric orientation of the three BB-appendages is suggested to be ideal to secure BBs to the cell cortex while ensuring BB organization and orientation (Allen, 1967, Iftode et al., 1996, Pitelka, 1961). However, the development, molecular regulators and functions of BB-appendage microtubules in creating connections with the cell cortex have not been closely studied.

**Figure 1:**
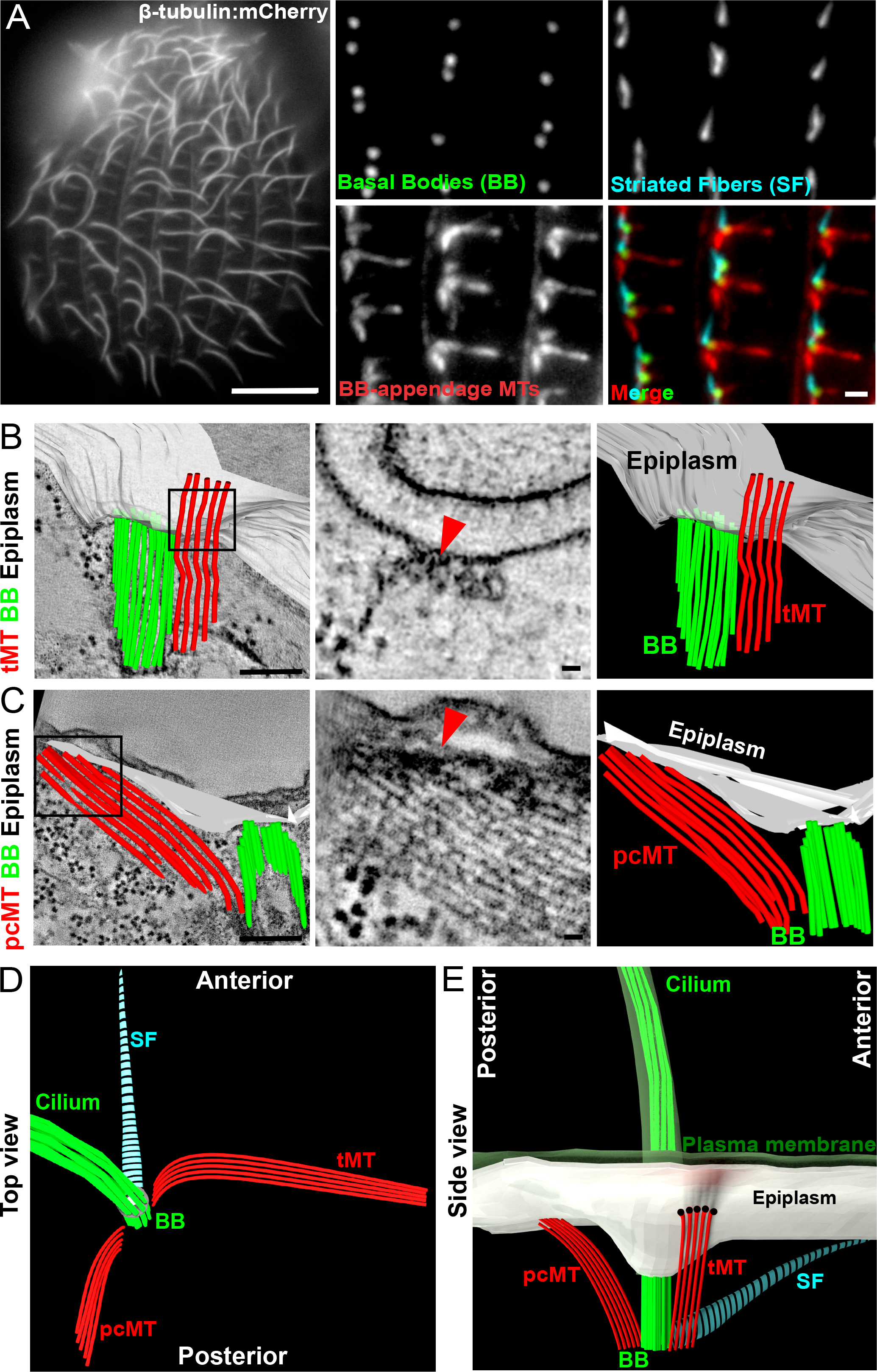
The BB-associated cortical cytoskeleton forms an organized pattern. (A) Left panel, the *Tetrahymena* cilia array (Btu2:mCherry). Scale bar, 10 µm. Right panels, BBs (Centrin, green), transverse (tMT) and post-ciliary (pcMT) microtubules (acetylated tubulin, red), and striated fibers (Bbc39:mCherry, cyan). Scale bar, 1 µm. (B) Left panel, 3D model of epiplasm (white), BB (green) and tMTs (red) projected on a tomographic slice. Boxed region highlights the tMT bundle running directly below the epiplasm. Scale bar, 200 nm. Middle panel, tomographic slice from the boxed region shows tMT connections with the cortical epiplasm (red arrowhead). Scale bar, 20 nm. Right panel, 3D model of the BB unit derived from EM tomographic reconstruction. (C) Left panel, 3D model of epiplasm (white), BB (green) and pcMTs (red) projected on a tomographic slice. Boxed region highlights pcMTs ending in the epiplasm. Scale bar, 200 nm. Middle panel, projected tomographic slices from the boxed region showing pcMTs ending in the cortical epiplasm (red arrowhead). Scale bar, 20 nm. Right panel, 3D model of the BB unit derived from EM tomographic reconstruction. (D) Top view model of a cilium, BB and BB-appendages. (E) Longitudinal view model of a cilium, BB and BB-appendages relative to the plasma membrane and epiplasm.

The cortical cytoskeleton is a network of cytoskeletal filaments lying just below the plasma membrane. This cortical cytoskeleton affects cell morphology, surface tension and elasticity while ensuring the positioning and orientation of BBs and their associated motile cilia (Discher et al., 1994, Zhang et al., 2017, Mahuzier et al., 2018, Herawati et al., 2016, Werner et al., 2011). During new BB assembly in ciliates, BBs are inserted through and attached to the cortical cytoskeleton (Aubusson-Fleury et al., 2013, Argetsinger, 1965, Allen, 1967, Pitelka, 1961). A major component of the ciliate cortical cytoskeleton, the epiplasm, is composed of intermediate filament-like proteins that form a dense fibrous network below the plasma membrane (Pitelka, 1961, Allen, 1967, Bouchard et al., 2001). Epc1 is the most abundant epiplasm component in *Tetrahymena* and *epc1*Δ mutants exhibit BB disorganization (Williams, 2004). Moreover, BB-appendage microtubules are physically linked to the epiplasm (Hufnagel, 1969, Allen, 1971, Allen, 1967, Iftode et al., 1996, Pitelka, 1961). These early studies postulated that BB-appendages are firmly attached to the epiplasm to maintain BB organization and orientation while resisting mechanical forces from beating cilia that would otherwise cause BBs to disorganize and disorient.

We investigated the development of BB-appendage microtubules following new BB assembly and the molecular events that promote BB cortical attachment, positioning and orientation within ciliary arrays. To understand the molecular underpinnings that govern the growth and attachment of BB-associated microtubules to the cortical cytoskeleton, we examined the effects of microtubule PTMs on BB organization. We identified microtubule glycylation to be required for normal BB-appendage microtubule elongation and BB cortical attachment, organization and orientation. Thus, microtubule glycylation has an important role in promoting BB and cilia positioning through associations with the cell cortex.

## RESULTS

### BB-associated microtubules contact the cortical cytoskeleton

*Tetrahymena* ciliary arrays are composed of hundreds of motile cilia (Fig. 1A). Cilia are nucleated and positioned by BBs, which have three appendage structures: the striated fiber (SF), the transverse microtubules (tMT), and the post-ciliary microtubules (pcMT). BBs and their associated appendage structures are organized into polarized rows and the direction that each appendage extends is uniform (Wloga and Frankel, 2012, Allen, 1967). We observe similar localization using immunofluorescence (Fig. 1A and S1A-D). Gaps in acetylated tMT labeling along the length of the tMT are attributed to masking of epitopes rather than a discontinuity within the microtubule structure. Thin section electron microscopy of *Paramecium* and *Tetrahymena* identified linkages between BB-appendage microtubules and the cortical epiplasm (Iftode et al., 1996, Allen, 1967, Pitelka, 1961). Using the improved 3D spatial resolution of electron tomography, we show that BB-appendage microtubules extend toward the cell cortex and contact the cortical epiplasm. tMTs extend from the anterior side of the BB toward the adjacent ciliary row (Fig. 1A,B; (Allen, 1967)). Electron dense linkages are detected at sites where the tMTs and the epiplasm are in close proximity (Figs. 1B, S1E and Movie1). Similarly, pcMTs emerge from the posterior face of BBs and extend posteriorly to contact the cortical epiplasm by two unique conformations (Figs. 1C, S1F and Movie2). First, the pcMT tips are embedded in an electron dense region of the epiplasm. The orientation and interface of tMT and pcMT attachments to the cortical epiplasm are unique. tMTs contact the cortical epiplasm by linkages between the bundle of microtubules running parallel to the epiplasm. Second, the pcMT ends interface with the cortical epiplasm by microtubule end-on linkages between the bundle of microtubules running perpendicular to the cortical epiplasm. The two conformations of pcMT interactions with the cortical epiplasm may reflect distinct mechanisms of BB anchoring and unique roles in resisting mechanical forces from beating cilia. Thus, BBs and BB-appendage microtubules are organized within ciliary arrays and are connected by distinct structural mechanisms to the cortical epiplasm (Fig. 1B-E and S1E, F).

### BB-appendage microtubules nucleate and elongate during new BB maturation

During *Tetrahymena* cellular growth, new BBs assemble into and expand the ciliary array (Nanney et al., 1975, Allen, 1969). New BBs form at the base of ciliated, mother BBs and mature as they migrate and separate in the cell’s anterior direction away from the mother BB (Fig. 2). The separation distance between mother and daughter BBs serves as a proxy for BB maturity (Pearson et al., 2009a, Perlman, 1973, Nanney et al., 1975). Once a new BB nucleates a cilium, it is considered mature. Because the maximum frequency of ciliated BBs (86%) is observed at a separation distance of 1.75 μm (Fig. S2A), BBs separated by less than 1.75 μm are on average still developing. To determine when maturing BBs acquire BB-appendage microtubules, the length of BB-appendage microtubules was measured relative to the BB separation distance (Fig. 2A-C). BBs and BB-appendage microtubules were labeled with endogenous mCherry (β-tubulin; Btu2:mCherry). The tMTs and pcMTs of new BBs initiate elongation concurrent with when new BBs can be resolved from mother BBs. As new BBs mature, the BB-appendage microtubules elongate to a maximum mean length of 1.4 µm and 0.5 µm for tMT and pcMTs, respectively. Most BBs are ciliated by a separation distance of 1.75 µm (Fig. S2A). This is the same point at which BB-appendage microtubule elongation begins to plateau, suggesting that tMTs and pcMTs complete elongation coincident with ciliogenesis. BB-associated microtubules are formed of linear bundles of microtubules (Fig. 2D-F). The pcMT and tMT bundles possess cross-linking densities between individual microtubules, thereby creating equal spacing between microtubules (Fig. 2E,F). Using EM tomography, we measured the total number of BB-appendage microtubules of new BBs relative to the distance separated from mother BBs (Fig. 2G and S2B). New BBs adjacent to their mother BBs do not possess BB-appendage microtubules. The number of microtubules in both tMT and pcMT bundles increases as new BBs mature reaching a maximum number of six microtubules. Thus, BB-appendage microtubule bundles are of full length and number before ciliogenesis. We hypothesize that BB-appendage microtubules achieve cortical attachments prior to ciliogenesis and prior to BBs experiencing ciliary forces. This may preserve the organization and orientation of BBs.

**Figure 2:**
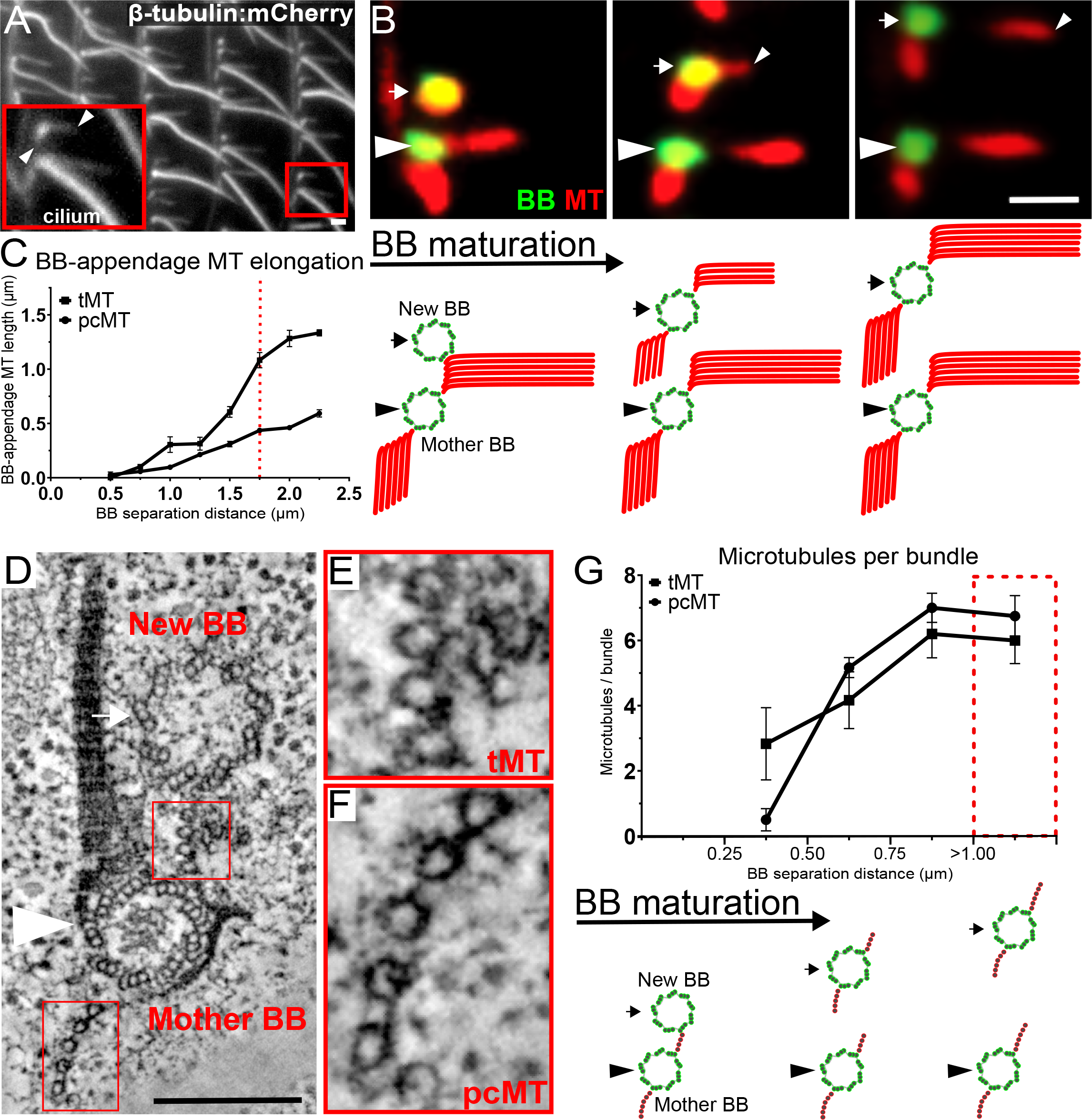
BB-appendage microtubule assembly during new BB maturation. (A) BBs with associated cilia and BB-appendage microtubules (Btu2:mCherry; grayscale). Inset shows a mother and new BB with arrowheads denoting the tMTs and pcMTs of the new BB. Scale bar, 1 µm. (B) Top panels, tMTs and pcMTs elongate coincident with BB maturation as determined by the separation distance between mother (arrowheads) and new BBs (arrows). BBs, Centrin, green; microtubules, acetylated-tubulin, red. Gap in tMTs is likely due to loss of antibody accessibility. Scale bar, 1 µm. Bottom panels, model of BB-appendage elongation with BB maturation. (C) tMTs and pcMTs elongate during BB maturation (n = 82 cells, 422 tMTs, 528 pcMTs). Dotted red line denotes BB separation distances when most BBs are ciliated (Fig. S2A). Quantification was performed on Btu2:mCherry cells (Fig. 2A). (D) EM tomographic slice of a mother and new BB pair. Scale bar, 200 nm. (E-F) Insets of the mother BB tMT and pcMT. Inset width 100 nm. (G) Top panel, the number of microtubules in tMT and pcMT bundles increases with BB maturation (n = 5 tomograms, 17 new BBs and corresponding tMT and pcMT bundles). Bottom panel, model of increasing BB-appendage microtubule number with BB maturation.

### Protein glycylation promotes BB organization and orientation

BBs and BB-appendage microtubules are dynamic structures that experience maturation, movement and ciliary forces. BB and BB-appendage microtubules carry several types of conserved tubulin PTMs that could be important in these processes. To determine the localizations of tubulin PTMs at BBs and BB-appendage microtubules, we utilized immunofluorescence imaging of PTM specific antibodies (Fig. 3). Glutamylated tubulin is enriched at BBs but is not detected at BB-appendage microtubules, consistent with our previous findings (Bayless et al., 2016). Glutamylation of tMTs and pcMTs was previously described (Wloga et al., 2008). However, after exploring multiple immunofluorescence preparations, glutamylated tubulin predominantly localized to BBs and we detected only a low level of glutamylated tubulin at tMTs and pcMTs. In contrast, acetylated and glycylated tubulin are both present at tMTs and pcMTs (Fig. 3; (Thazhath et al., 2002, Tassin et al., 2015, Iftode et al., 2000)). Antibodies recognizing microtubule monoglycylation and polyglycylation mark both tMTs and pcMTs. These observations indicate that acetylation and glycylation are enriched on the BB-appendage microtubules.

**Figure 3:**
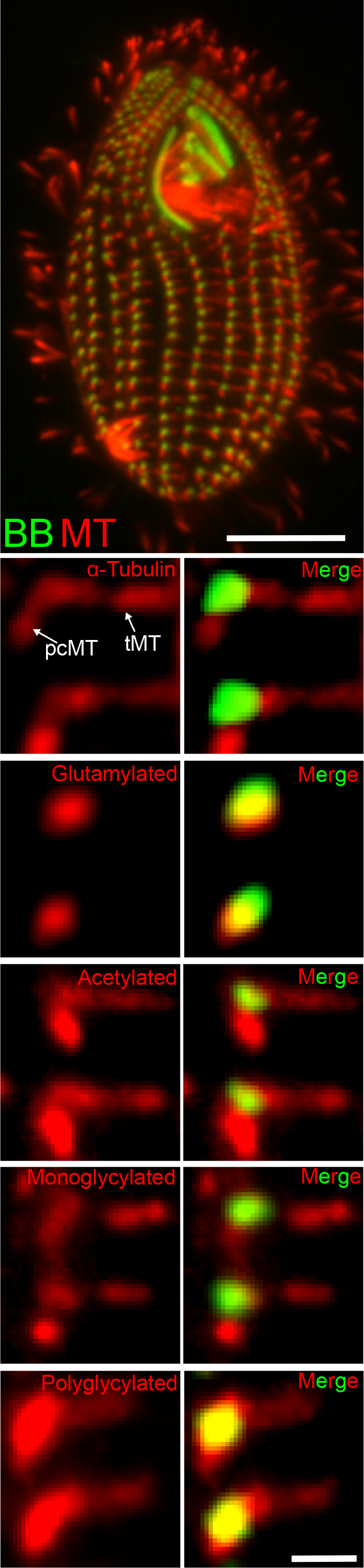
BB-appendage microtubules are enriched for acetylation and glycylation microtubule post-translational modifications. *Tetrahymena* BBs (Centrin, green) and microtubules (acetylated-tubulin, red). Scale bar, 10 µm. Bottom panels, localization of tubulin and tubulin PTMs (red) relative to the BBs (Centrin, green). White arrows denote the locations of tMTs and pcMTs. Scale bar, 1 µm.

To test whether microtubule acetylation promotes BB organization, we visualized BB organization in mutants that lack microtubule acetylation. Both *mec17*Δ (acetyltransferase knockout) and *atu1-K40R* (non-acetylatable α-tubulin mutant) mutants eliminate detectable microtubule acetylation (Gaertig et al., 1995, Akella et al., 2010). However, both mutant cell lines exhibit normal BB organization (Fig. S2C-D).

Glycylation modifies tubulin C-terminal tails and is present at BB-appendage microtubules (Fig. 3). Ttll3A,B,C,D,E,F include all *Tetrahymena* proteins with homology to the Ttll3 and Ttll8 monoglycylases. To test whether microtubule glycylation promotes BB organization, we examined the *ttll3aΔ,bΔ,cΔ,dΔ,eΔ,fΔ* glycylase mutants (hereafter *ttll3Δ*; (Wloga et al., 2009)). After *TTLL3* knockout, cells lose microtubule glycylation (Fig. 4A). Prior studies of these mutants reported slow cell multiplication rates and shorter cilia but did not note BB organization defects (Wloga et al., 2009). However, detailed analyses showed *ttll3Δ* cells to exhibit BB disorganization. BB disorganization was defined as cells with greater than three BBs that were mislocalized from the axis of a ciliary row by at least one μm. BBs are organized correctly in *ttll3Δ* cells immediately following *TTLL3* knockout (Day 0). This is likely because the progeny cells have remaining parental Ttll3 protein (Figs. 4B and S3A). The BB disorganization increases and peaks two days after *TTLL3* knockout, when 94% of *ttll3Δ* cells possess disorganized BBs. The severity of BB disorganization in the *ttll3Δ* cells ranges from 3 to 54 disorganized BBs per cell (Fig. 4B and S3A). In summary, Ttll3, likely through microtubule glycylation, is important for organizing BBs within ciliary arrays.

**Figure 4:**
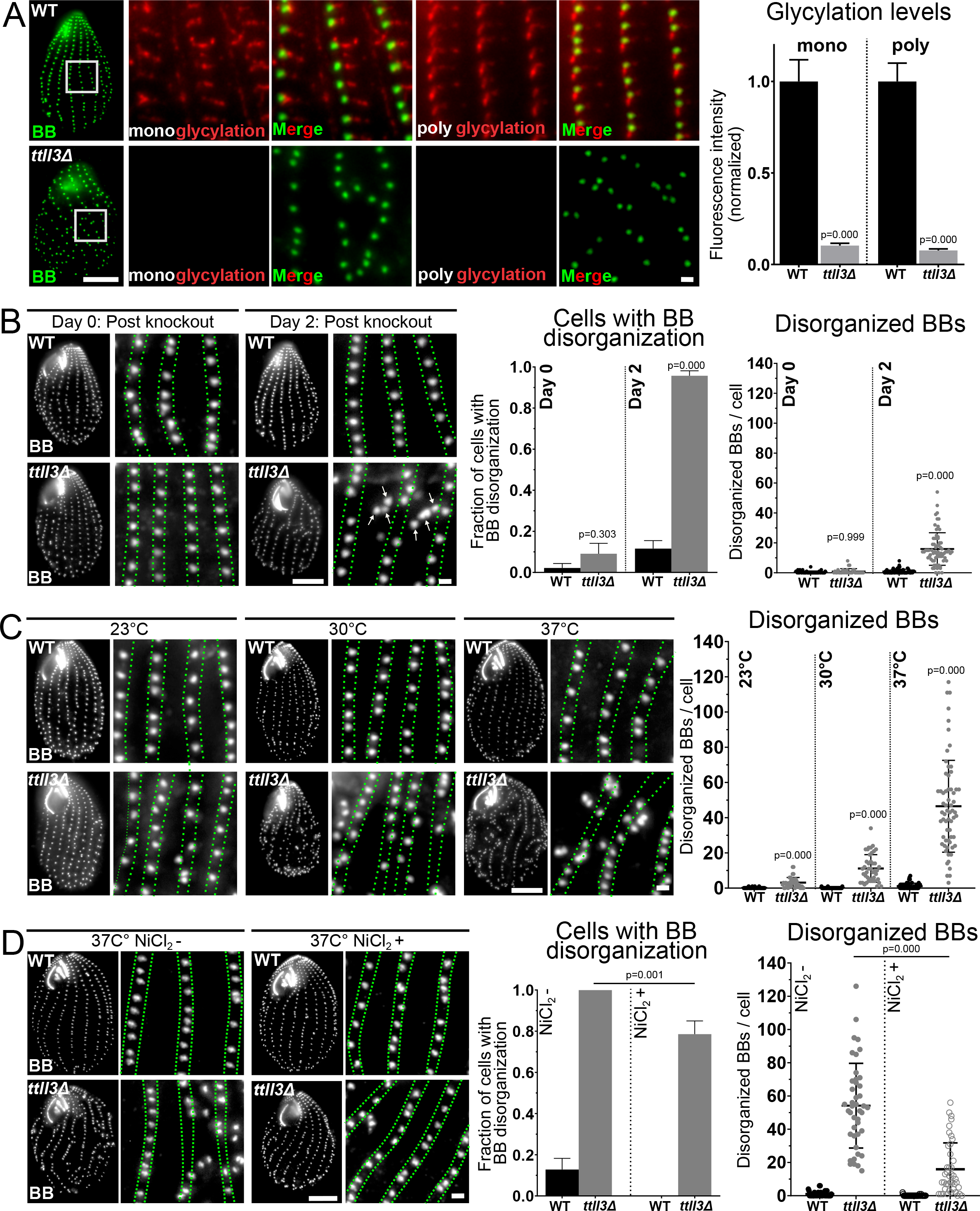
Microtubule glycylation promotes BB organization and appendage microtubule length. (A) Left panels, WT and *ttll3Δ* BBs (Centrin, green). Scale bar, 10 µm. Insets show loss of mono- and poly-glycylation in *ttll3Δ* cells two days post knockout (BB, Centrin, green; BB-appendage microtubules, glycylation, red). Scale bar, 1 µm. Right panel, *ttll3Δ* cells have reduced glycylation (WT monoglycylation: n = 42 cells, *ttll3Δ* monoglycylation: n = 33 cells, WT polyglycylation: n = 34 cells, *ttll3Δ* polyglycyation: n = 42 cells). (B) Left panels, WT and *ttll3Δ* cells cultured 0 and 2 days post knockout (BB, Centrin, grayscale). Green lines denote boundaries of BB rows. Arrows denote disorganized BBs. Scale bars, 10 and 1 (inset) µm. Right panels, increased fraction of *ttll3Δ* cells at 30ºC have BB disorganization on Day 2 (WT day 0: n = 46 cells, *ttll3Δ* day 0: n = 33 cells, WT day 2: n = 69 cells, *ttll3Δ* day 2: n = 70 cells). *ttll3Δ* cells at 30ºC have more disorganized BBs per cell on Day 2 (WT day 0: n= 46 cells, *ttll3Δ* day 0: n = 33 cells, WT day 2: n = 69 cells, *ttll3Δ* day 2: n = 70 cells). (C) Left panels, WT and *ttll3Δ* cells cultured at 23ºC, 30ºC and 37ºC (BB, Centrin, grayscale). Green lines denote boundaries of BB rows. Scale bars, 10 and 1 (inset) µm. Right panels, *ttll3Δ* cells have increasing BB disorganization with increasing temperature (WT 23ºC: n = 35 cells, *ttll3Δ* 23ºC: n = 42 cells, WT 30ºC: n = 40 cells, *ttll3Δ* 30ºC: n=43 cells, WT 37ºC: n = 54 cells, *ttll3Δ* 37ºC: n = 63 cells). (D) Left panels, WT and *ttll3Δ* cells cultured at 37ºC in the presence or absence of NiCl_2_ (Centrin, grayscale). Green lines denote boundaries of BB rows. Scale bars, 10 and 1 (inset) µm. Right panels, NiCl_2_ treatment of *ttll3Δ* cells partially reduces BB disorganization (WT NiCl_2_-: n = 40 cells, *ttll3Δ* NiCl_2_-: n = 43 cells, WT NiCl_2_+: n = 38 cells, *ttll3Δ* NiCl_2_+: n = 42 cells).

Motile cilia are oriented along the cell’s anterior-posterior axis (Fig. 1). To determine whether BBs in *ttll3Δ* cells have orientation defects, we quantified the orientation of BBs in *ttll3Δ* cells either inside (organized) or outside (disorganized) of ciliary rows, using the SF BB-appendage as a BB polarity marker (Fig. S3B). Circular statistical analyses were used to assess BB orientation. An R-value of 1.00 indicates perfectly orientated BBs while an R-value of 0.00 indicates random BB orientation. *ttll3Δ* BBs inside ciliary rows (organized) were 6% less oriented (R-value = 0.93) than WT BBs (R-value = 0.99). Additionally, disorganized *ttll3Δ* BBs were less oriented than organized *ttll3Δ* BBs (R-value = 0.75). Disorganized BBs were not detected in WT cells. Thus, microtubule glycylation promotes the organization and orientation of BBs.

### Glycylation stabilizes BB organization against forces from ciliary beating

BBs remain organized and oriented while anchoring motile cilia that produce mechanical forces. Ciliary beat frequency and ciliary forces transmitted to BBs can be manipulated in *Tetrahymena* by increasing temperature or media viscosity (Galati et al., 2014, Pearson et al., 2009b, Goto et al., 1982). BB disorganization in *ttll3Δ* cells was found to recover at lower temperatures (23ºC; Fig. 4C and S3A). Because low temperatures reduce ciliary beating, this result suggested that the BB organization phenotypes in *ttll3Δ* cells are responsive to ciliary forces. After prolonged culture at 23ºC, *ttll3Δ* cells were shifted to 30ºC or 37ºC for 24 hrs (Fig. 4C). 93% of *ttll3Δ* cells shifted to 30ºC exhibited an intermediate level of BB disorganization. Finally, 100% of *ttll3Δ* cells at 37ºC had severe BB disorganization. The severe BB disorganization in *ttll3Δ* cells at 37ºC was reduced by decreasing ciliary beating with NiCl_2_, an inhibitor of ciliary inner dynein arms (Fig. 4D; (Larsen and Satir, 1991)). *Tetrahymena* cells divide more rapidly at elevated temperatures and new BB biogenesis and organization is maintained through these rapid cell divisions (Frankel, 1962, Frankel and Nelsen, 2001, Hallberg and Hallberg, 1983). To further test whether the BB disorganization in *ttll3Δ* cells at elevated temperature is a result of increased ciliary forces and not aberrant maturation from increased cell division and BB assembly rates, we measured the cell division rates of *ttll3Δ* cells with and without NiCl_2_ (Fig. S3C). No difference was observed, suggesting that a rapid rate of BB assembly and maturation is not responsible for BB disorganization in *ttll3Δ* cells. Surprisingly, BB disorganization was not elevated in *ttll3Δ* cells exposed to increased media viscosity which also elevates mechanical forces on cilia (Fig. S3D). This suggests that glycylation protects the organized ciliary array from temperature-induced high frequency ciliary forces but does not play a role in resisting viscosity-induced elevated ciliary load. This is in contrast to strains mutated in the SF BB-appendage protein DisA that are sensitive to increases in ciliary load, suggesting that different appendage structures have distinct roles at the BB in counteracting the forces produced by ciliary beating (Galati et al., 2014, Jerka-Dziadosz et al., 1995). In summary, tubulin glycylation is important to maintain BB organization when confronted with rapid ciliary beating.

Our studies suggest that tubulin glycylation is important for maintaining BB organization and orientation even in the face of ciliary forces. *Tetrahymena* cells are asymmetrically shaped with dorsal-ventral and anterior-posterior axes and we hypothesized that specific regions of the cell may be more sensitive to ciliary forces. To test whether specific regions of the *Tetrahymena* cell geometry are more sensitive to BB disorganization, we quantified the position of BB disorganization. BB disorganization is most frequent in the medial and posterior regions along the dorsal axis of the cell (Fig. S4A). This suggests that BBs and BB-appendage microtubules at the dorsal and medial regions of *Tetrahymena* cells are subjected to greater physical forces from beating cilia.

The BB disorganization and disorientation observed in *ttll3Δ* cells suggests that tubulin glycylation might promote attachments between BBs and the cell cortex. To determine whether BBs in *ttll3Δ* cells were cortically detached, we visualized BBs relative to the cell cortex. Cortically detached BBs were defined as BBs that were internally separated from the cell cortex by greater than one μm. 24% of *ttll3Δ* cells at 30ºC possessed BBs that were detached from the cell cortex compared to 5% for WT cells (Fig. 5A and S4B). Cortically detached BBs were observed as either individual BBs that were 1-3 μm internal to the cell cortex or clustered BBs within the cell’s interior (Fig. S4B). In summary, reduced microtubule glycylation results in BB detachment from the cell cortex.

**Figure 5:**
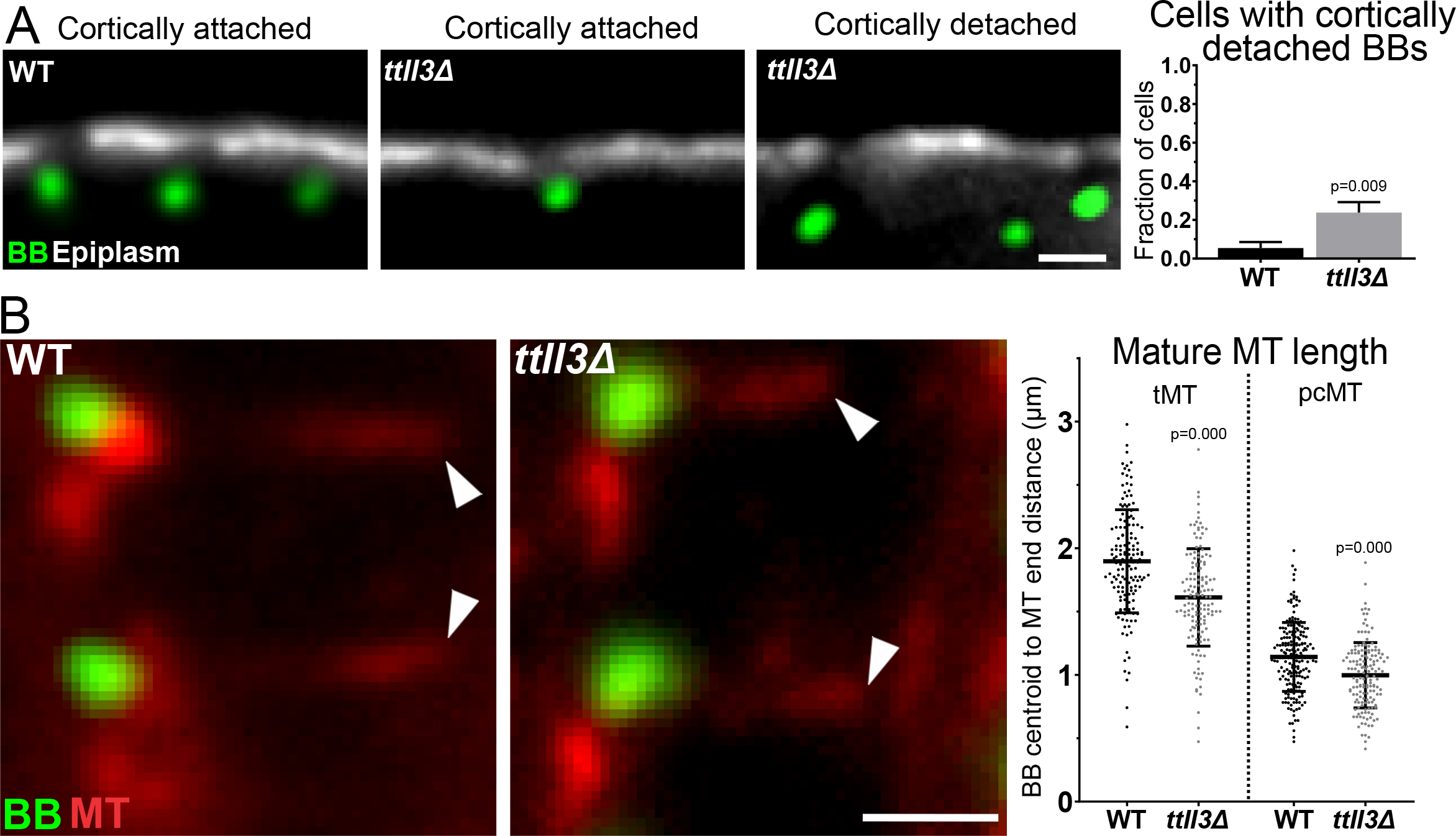
Glycylation promotes BB cortical attachment and BB-appendage microtubule length. (A) Left panels, longitudinal sections through BBs (Centrin, green) and epiplasm (α-Epc1, grayscale) in WT and *ttll3Δ* cells. Right panel, *ttll3Δ* cells have more cortically detached BBs at 30ºC (WT: n = 55, *ttll3Δ*: n = 63). Scale bar, 1 µm. (B) BBs (Centrin, green) and BB-appendage microtubules (α-tubulin, red) in WT and *ttll3Δ* cells. Right panel, tMTs and pcMTs of mature BBs are shorter in *ttll3Δ* cells (WT: n = 67 cells, 156 tMTs, 194 pcMTs; *ttll3Δ*: n = 75 cells, 141 tMTs, 155 pcMTs).

### Glycylation promotes BB-appendage length

The reductions in BB attachment, organization and orientation in *ttll3Δ* mutant cells suggest that glycylation promotes BB positioning by promoting attachment to the cortical epiplasm. BB-appendage microtubules are near their fully elongated state by the time of ciliogenesis when daughter BBs are separated from mother BBs by 1.75 μm (Fig. 2). We hypothesized that BB-appendage microtubule elongation ensures BB attachment to the cell cortex and next explored whether glycylation could promote BB attachment to the cell cortex by promoting the normal lengths of BB-appendage microtubules. To quantify the length of tMTs and pcMTs in WT and *ttll3Δ* cells, antibodies specific to BBs and α-tubulin were used to mark these structures and the distances between the BB centroid to the ends of the tMTs and pcMTs were measured. Because the maximum frequency of ciliated BBs (86%) is observed at a separation distance of 1.75 μm (Fig. S2A), BBs separated by less than 1.75 μm are on average still developing. tMTs and pcMTs associated with both developing and mature BBs are shorter in *ttll3Δ* cells than WT controls (Figs. 5B and S4C). Therefore, microtubule glycylation accelerates the rate of new BB-appendage microtubule elongation and promotes a longer length for these microtubules at maturity.

### BB organization and attachment to the cortical epiplasm

BB-appendage microtubules contact the cell cortex by attaching to the cortical epiplasm that lies just below the plasma membrane (Figs. 1 and S1). We hypothesized that disrupting the cortical epiplasm would therefore produce analogous BB positioning defects to those observed in *ttll3Δ* mutant cells. Indeed, BB organization and cortical attachment is disrupted in *epc1Δ* epiplasm mutant cells (Fig. 6; (Williams, 2004)). Like *ttll3Δ* cells, Williams reported that BB organization returned to normal within approximately 20 generations after *EPC1* knockout. To test if Epc1 has a role in maintaining BB organization when ciliary forces are increased, we measured BB disorganization in *epc1Δ* cells cultured at elevated temperatures. At 23ºC, *epc1Δ* cells did not exhibit detectable BB disorganization. However, upon shifting *epc1Δ* cells to 30ºC and 37ºC, BB disorganization increased (Fig. 6B). Thus, the epiplasm is part of the system that maintains BB organization in the face of ciliary forces.

**Figure 6:**
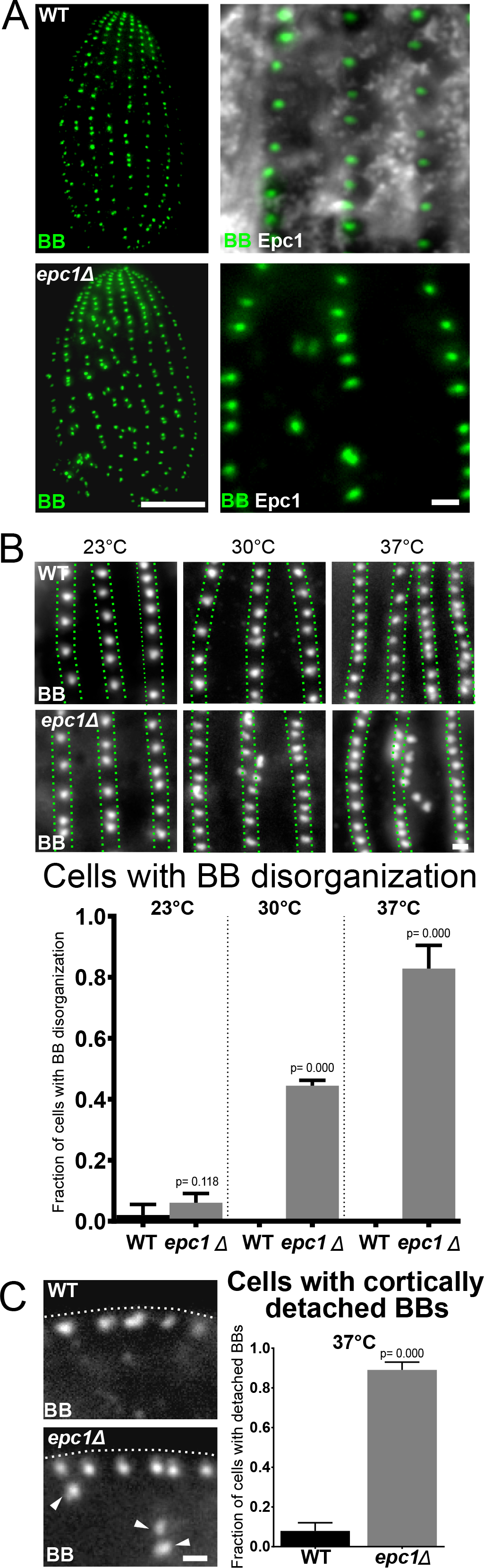
Cortical epiplasm is required for BB organization and cortical attachment. (A) the dorsal side (non-oral apparatus) of WT and *epc1Δ* cells labeled for BBs (Centrin, green). Scale bar, 10 µm. Inset, BBs (Centrin, green) and epiplasm (α-Epc1, grayscale) in WT and *epc1Δ* cells. Scale bar 1 µm. *epc1Δ* cells have disorganized BBs. (B) Top panels, BBs (Centrin, grayscale) labeled in WT and *epc1Δ* cells grown at 23ºC, 30ºC and 37ºC for 24 hrs. Scale bar, 1 µm. Bottom panels, elevated temperature increases BB disorganization in *epc1Δ* cells (WT 23ºC: n = 99 cells, *epc1Δ* 23ºC: n = 99 cells, WT 30ºC: n = 99 cells, *epc1Δ* 30ºC: n = 99 cells, WT 37ºC: n = 99 cells, *epc1Δ* 37ºC: n = 99 cells). (C) Left panels, longitudinal section through BBs (Centrin, grayscale) at the cell cortex in WT and *epc1Δ* cells. White arrowheads denote BBs that are detached from the cell cortex. White line denotes the location of cell cortex as determined by the cellular fluorescence background. Scale bar, 1 µm. Right panels, *epc1Δ* cells have more cortically detached BBs (WT 37ºC: n= 63 cells, *epc1Δ* 37ºC: n=64 cells).

Epc1’s role in organizing BBs and possibly resisting forces from cilia beating suggested that the epiplasm is important to anchor BBs to the cell cortex. To test this, we measured the cortical attachment of BBs in *epc1Δ* cells cultured at 37ºC when BBs are severely disorganized. BB attachment to the cell cortex was determined by immunofluorescence localization of BBs relative to differential interference contrast (DIC) imaging of the cell cortex. 90% of *epc1Δ* cells exhibited cortically detached BBs (Fig. 6C). Together, this suggests that the epiplasm and glycylated BB-appendage microtubules are both important in cortically attaching and organizing and cortically attaching BBs.

## DISCUSSION

Cilia and BBs are attached to the cell cortex and organized and oriented within polarized arrays to produce effective hydrodynamic flow. BB-associated microtubules are vital to ensure this organization (Werner et al., 2011, Herawati et al., 2016, Kunimoto et al., 2012). We show that *Tetrahymena* BB-appendage microtubules nucleate and elongate from new BBs coincident with BB maturation so that they are poised for cortical attachment by the time of ciliogenesis. Ttll3 glycylases, likely through microtubule glycylation, promote efficient elongation of BB-appendage microtubules and maintain BB organization and orientation. Further, both BB-appendage microtubules and the cortical cytoskeleton that they attach to promote BB attachment and organization. This study illuminates the importance of post-translationally modified BB-associated microtubules in connecting BBs to the cortical cytoskeleton to resist forces from beating cilia.

### BB-appendage microtubule length for cortical attachment of BBs

Zebrafish Ttll3 knockdown reduces polarized ciliary beating suggesting that tubulin glycylation is important for either cilia beating and/or BB orientation (Pathak et al., 2011). In ciliates, BB-appendage microtubules are post-translationally modified, however, it was not known whether and how PTMs promote BB organization and orientation (Fig. 3; (Wloga et al., 2009, Callen et al., 1994, Tassin et al., 2015, Gaertig et al., 1995)). Here, we provide the first evidence that glycylation of *Tetrahymena* BB-appendage microtubules promotes BB organization, orientation and cortical attachment.

Supporting a model where BB-appendage microtubule length contributes to the stabilization of BB organization, loss of microtubule glycylation reduces the length of BB-appendage microtubules and disorganizes BBs. Given that BB-appendage microtubules in *ttll3Δ* cells are shorter during BB maturation, microtubule glycylation likely affects elongation of these microtubules. This is consistent with tubulin glycylation’s role in promoting both the elongation of *Tetrahymena* cilia and in the maintenance of mouse ependymal cell and retinal cell cilia length (Wloga et al., 2009, Bosch Grau et al., 2017, Gadadhar et al., 2017b). We suggest that the length of BB-appendage microtubules is important for cortical contacts to anchor BBs and to reduce BB mobility resulting from ciliary beating forces.

Upon cilia formation and beating, BBs experience mechanical forces from undulating cilia (Galati et al., 2014, Pearson et al., 2009b, Bayless et al., 2016). Consistent with this, when BB-appendage microtubules or the cortical cytoskeleton are disrupted, BB disorganization is increased when ciliary beating is elevated by increasing temperature. Reduced ciliary beating even at high temperatures reduces this BB disorganization. Surprisingly, *ttll3Δ* mutants are not sensitive to increased ciliary load when cells are subjected to increased viscosity. This contrasts the *disA-1* SF mutant that is sensitive to both elevated temperature and viscosity (Galati et al., 2014). Elevated viscosity is reported both to alter ciliary and flagellar waveform and to slow ciliary beat frequency (Sleigh, 1966, Rikmenspoel, 1971, Minoura and Kamiya, 1995, Machemer, 1972). Ciliary dynein arm mutations alter the ciliary beat pattern and frequency when exposed to higher viscosity media, underscoring the importance of ciliary structure for cellular response to high viscosity conditions (Wilson et al., 2015). Because *ttll3Δ* mutants harbor shorter cilia, we suspect that ciliary beat patterns are affected in *ttll3Δ* mutant cells at high viscosity to somehow reduce the forces transduced to BBs. Regardless, BB-appendage microtubule glycylation promotes their elongation and attachment of BBs to the cell cortex to resist mechanical forces from ciliary beating.

### BB-appendage microtubule glycylation for cortical attachment of BBs

As BB-appendage microtubule bundles elongate, the microtubule surface area juxtaposed to the epiplasm is increased (Fig. 1D, E). We hypothesize that this increases cortical contacts to anchor BBs and to reduce BB mobility that result from forces of ciliary beating. Tubulin glycylation modifies the surface of microtubules by creating a layer of flexible, uncharged residues (Wall et al., 2016, Gadadhar et al., 2017a). Thus, in addition to promoting BB-appendage microtubule growth, glycylation may also control the accessibility of the charged residues on tubulin’s C-terminal tail. This may alter protein associations with these microtubules and promote BB organization by increasing the affinity of BB-appendage microtubules for the epiplasm. Thin section EM studies and our electron tomography data show electron dense connections between the epiplasm and tMTs (Figs. 1B and S1E; (Allen, 1967, Iftode et al., 1996)). This would suggest that these microtubules do not need to contact the epiplasm directly but are physically linked through associated proteins that remain to be discovered. The ends of the pcMT are embedded within the epiplasm. This interaction could be direct or rely on additional proteins. Given that both tMTs and pcMTs are grossly modified in similar ways, how their interactions with the cell cortex are differentially regulated is an interesting question. There may be differences in glycylation chain length, or in specific residues on the C-terminal tails that receive modifications.

### Recovery of BB organization and orientation

Despite the loss of BB-appendage microtubule glycylation, the BB disorganization phenotype in *ttll3Δ* cells recover several days after initiation of *TTLL3* knockout (Fig. S3A). BB disorganization can almost completely recover if grown for several weeks at 23ºC, where ciliary forces are reduced (Fig. 4C). Thus, *Tetrahymena* cells compensate for the loss of microtubule glycylation and recover from BB disorganization. However, the reorganized BBs in *ttll3Δ* remain sensitive to high temperature. We suggest that the temperature sensitivity results from the elevated ciliary forces that are associated with temperature-induced high frequency cilia beating. Similar to the compensation observed in *ttll3Δ* cells, BB disorganization in *epc1Δ* cells recovers after extended growth at low temperature (Williams, 2004). Thus, *Tetrahymena* cells compensate for cortical morphology defects that arise from disruption of tubulin glycylation and the cortical cytoskeleton that they attach to. We did not observe changes to the levels or localization of Epc1 when Ttll3 was lost, nor in the levels or localization of BB-associated microtubule acetylation or glycylation when Epc1 was lost. Tubulin glutamylation increases in *ttll3Δ* cells (Wloga *et al.*, 2009), however this increase is not due to the recruitment of glutamylation to BB-appendage microtubules (Fig. S4E). It is possible that the compensation is mediated through other cortical cytoskeletal proteins (Epiplasmin and K-antigen proteins; (Williams et al., 1995, Williams et al., 1990, Honts and Williams, 2003)). The localization of these cortical proteins is consistent with the sites of BB-appendage microtubule association with the epiplasm. In summary, our studies suggest that multiple pathways promote effective BB organization and orientation.

### Glycylation of non-tubulin targets

Tubulin is not the only protein that is glycylated by Ttll enzymes. Nucleosome Assembly Protein 1 (NAP-1) is glycylated by Ttll10 in addition to other nucleosome proteins that are glycylated by Ttll8 in *Drosophila* (Ikegami et al., 2008, Rogowski et al., 2009). However, there is no indication that nucleosome component glycylation regulates BB organization. Moreover, *Giardia* TTLL3 polygylcylates C-terminal glutamate residues on 14-3-3 (epsilon-like) that regulates actin organization (Lalle et al., 2011, Lalle et al., 2006, Krtkova et al., 2017). A *Tetrahymena* 14-3-3, Ftt18, was previously reported at BBs and pcMTs (Kilburn et al., 2007). However, it is not known whether *Tetrahymena* Ftt18 is glycylated. In fact, tubulin and the Hsp70 family protein, Pgp1, are the major glycylated *Tetrahymena* proteins (Xie et al., 2007). Pgp1 is an essential ER-associated chaperone protein that responds to protein folding stress including those at elevated temperatures. Interestingly, Pgp1 glycylation likely occurs within the ER by Ttll enzymes with ER signal sequences (Xie et al., 2007). Importantly, the glycylated tubulin antibodies used in our study do not recognize the ER. Taken together, while we cannot exclude contributions from other glycylated proteins, the observed phenotypes in *ttll3Δ* cells are consistent with tubulin as the primary glycylated substrate for BB positioning and attachment to the cell cortex.

### BB-appendages and cortical attachments for cell morphology

BB-associated microtubules provide a structural framework for establishing and maintaining cell shape (Pitelka, 1961, Allen, 1967, Tilney et al., 1966, Tilney and Porter, 1965). Consistent with this, *Tetrahymena ttll3Δ* cells are rounder and more variable in shape than WT (Figs. 4, S3 and S4). We suggest that reduced cortical attachments of the BB and BB-appendage microtubules decreases their ability to effectively serve as a structural scaffold. Moreover, BB rows in *ttll3Δ* cells are commonly skewed relative to the geometry of the cell thereby creating ciliary rows that spiral around the cell (Fig. 4). This is reminiscent of the *twi1-1* (*scr1-1*) mutant that exhibits twisting of BB rows in a temperature sensitive manner (Frankel, 2008). Additionally, we find that the medial and posterior regions along the dorsal axis of *ttll3Δ* cells have more disorganized BBs. We suggest that BBs within these regions of *Tetrahymena* cells experience greater ciliary forces. Tubulin glycylation appears to be important for anchoring to resist these differential forces experienced across the cell’s geometry. In summary, tubulin glycylation of BB-appendage microtubules is important for both local BB organization and orientation and global cellular morphology. Glycylated BB-appendage microtubules, in combination with the cortical cytoskeleton, provide a structural framework to maintain cell shape.

## Abbreviations

BB: basal bodies
SF: striated fiber
tMT: transverse microtubule
pcMT: post-ciliary microtubule
MT: microtubule
PTM: post-translational modification

## SUPPLEMENTAL FIGURE LEGENDS

**Figure S1: *Tetrahymena* cortical cytoskeleton.** (A) Left and top panels, Epc1:mCherry (grayscale) labeled cell with BBs (Centrin, green) and BB-appendage microtubules (polyglycylated tubulin, red). Scale bars, 10 and 1 (inset) µm. (B) Single Z-plane images of longitudinal sections through BB units showing the relative proximity of tMTs and the epiplasm (Centrin, green; acetylated-tubulin, red; Epc1:mCherry, grayscale). Scale bar, 1 µm. (C) Single Z-plane images of longitudinal sections through BBs show the relative proximity of pcMTs to the epiplasm (Centrin, green; polyglycylation, red; Epc1:mCherry, grayscale). Scale bar, 1 µm. (D) Left panel, BBs (Centrin, green), cilia and BB-appendage microtubules (glutamylated tubulin, red), and epiplasm (Epc1:mCherry, grayscale). Scale bar, 10µm. Right panels, single Z-plane images of a longitudinal section through BB units. Scale bar, 1 µm. (E) Left panels, 3D models of epiplasm (white), BB (green) and tMTs (red) projected on tomographic slices. The boxed regions highlight tMT bundles running directly below the epiplasm. Scale bar, 200 nm. Middle panels, tomographic slices from the boxed regions show tMT connections with the cortical epiplasm (red arrowheads). Scale bar, 20 nm. Right panels, 3D model of the BB units derived from EM tomographic reconstructions. (F) Left panels, 3D models of epiplasm (white), BB (green) and pcMTs (red) projected on tomographic slices. Boxed regions highlight pcMT bundles running directly below the epiplasm. Scale bar, 200 nm. Middle panels, tomographic slices from the boxed regions show pcMT-Epiplasm interfaces (red arrowheads). Scale bar, 20 nm. Right panels, 3D models of the BB units derived from EM tomographic reconstructions.

**Figure S2: Microtubule acetylation is not required for BB organization.** (A) Left panel, *Tetrahymena* cilia (glutamylated-tubulin, red) and BBs (Centrin, green). Scale bar, 1 µm. Right panel, the frequency of ciliated new BBs increases with separation distance from mother BBs (n = 52 cells, 648 ciliated BB pairs). (B) Quantification of the number of microtubules in tMT and pcMT bundles of mature BBs. (C) Left panels, BBs (Centrin, green) in WT and *mec17Δ* cells. Scale bar, 10µm. Middle panels, insets of WT and *mec17Δ* BBs (Centrin, green) and BB-appendage microtubules (acetylated-tubulin red). Scale bar, 1 µm. Right panel, BB disorganization does not increase in *mec17Δ* cells lacking acetylated-tubulin (WT: n = 55 cells, *mec17Δ*: n = 51 cells). (D) Left panels, WT and *K40R* BBs (Centrin, green). Scale bar, 10µm. Middle panels, insets of WT and *K40R* BBs (Centrin, green) and BB-appendage microtubules (acetylated-tubulin, red). Scale bar, 1 µm. Right panel, BB disorganization does not increase in *K40R* cells (WT: n = 55 cells, *K40RΔ*: n = 48 cells).

**Figure S3: Glycylation promotes BB organization against ciliary forces.** (A) Left panels, WT and *ttll3Δ* BBs (Centrin, grayscale) in cells grown 0, 1, 2, 3, 4 and 5 days post knockout at 30ºC. Scale bars, 10 and 1 (inset) µm. Green lines denote boundaries of BB rows. Arrows indicate disorganized BBs. Top right panel, fraction of cells with BB disorganization decreases after Day 2 in *ttll3Δ* cells (WT day 0: n = 46 cells, *ttll3Δ* day 0: n = 33, WT day 1: n = 30, *ttll3Δ* day 1: n = 30, WT day 2: n = 69, *ttll3Δ* day 2: n = 70, WT day 3: n = 30, *ttll3Δ* day 3: n = 30, WT day 4: n = 30, *ttll3Δ* day 4: n = 30, WT day 5: n = 30, *ttll3Δ* day 5: n = 30). Bottom right panel, the number of disorganized BBs per *ttll3Δ* cell decreases after Day 2 (WT day 0: n = 46, *ttll3Δ* day 0: n = 33, WT day 1: n = 30, *ttll3Δ* day 1: n = 30, WT day 2: n = 69, *ttll3Δ* day 2: n = 70, WT day 3: n = 30, *ttll3Δ* day 3: n = 30, WT day 4: n = 30, *ttll3Δ* day 4: n = 30, WT day 5: n = 30, *ttll3Δ* day 5: n = 30). B) Left panels, BB (Centrin, green) and SF (α-striated fiber, magenta) orientation in WT and *ttll3Δ* cells. Middle panel, organized and disorganized *ttll3Δ* BBs are disoriented (WT: n = 58 cells, WT organized BBs and SFs = 1235, *ttll3Δ* cells: n = 64 cells, *ttll3Δ* organized and disorganized BBs and SFs = 1567). Right panel, arrow graphs represent the frequency of angular deviation from the cell anterior-posterior axis observed in disoriented BBs. (C) NiCl_2_ does not affect the number of WT or *ttll3Δ* cell divisions in 24 hrs (WT NiCl_2_-: n = 3, WT NiCl_2_+: n = 3, *ttll3Δ* NiCl_2_-: n = 3, *ttll3Δ* NiCl_2_+: n = 3). (D) Left panels, WT and *ttll3Δ* BBs (Centrin, grayscale) in cells cultured at 30ºC in standard and viscous media. Scale bars, 10 and 1 (inset) µm. Green lines denote BB rows. Right panel, BB disorganization does not increase in *ttll3Δ* cells in high viscosity media (WT standard viscosity: n = 46 cells, *ttll3Δ* standard viscosity: n = 62 cells, WT high viscosity: n = 49 cells, *ttll3Δ* high viscosity: n = 62 cells).

**Figure S4: BB disorganization and slower BB-appendage microtubule elongation in ttll3**Δ **cells.** (A) BB (Centrin, grayscale) organization along the ventral and dorsal axis of WT and *ttll3Δ* cells. The medial and posterior regions of the dorsal side of *ttll3Δ* cells exhibited increased BB disorganization. Green lines denote boundaries between the anterior quartile, medial half, and posterior quartile. Scale bar, 10 µm. (B) Longitudinal view of BBs (Centrin, green) and SFs (α-SF, magenta) relative to the cell outline. Detached BBs are either near the cortex or in internal clusters. Scale bars, 10 and 1 (inset) µm. Arrows denote detached BBs. (C) Left panels, BBs (Centrin, green) and BB-appendage microtubules (α-tubulin, red) in WT and *ttll3Δ* cells during BB maturation. Arrowheads denote mother BBs and arrows denote new BBs. Scale bar, 1 µm. Right panels, elongating tMT and pcMT lengths are shorter in *ttll3Δ* cells (WT: n = 67 cells, 258 tMTs, 331 pcMTs; *ttll3Δ*: n = 75 cells, 379 tMTs, 396 pcMTs). Asterisks denote p value <0.05. (D) WT and *ttll3Δ* cells cultured for >30 days at 23ºC, shifted to 30ºC for 24 hrs and stained for BBs (Centrin, green) and glutamylated tubulin (GT335, red). Arrowheads denote glutamylated tubulin stained cilia. Scale bars, 10 and 1 (inset) µm.

## MATERIALS AND METHODS

### *Tetrahymena* strains

*Tetrahymena thermophila* cells (CU428, SB1969 and B2086) were obtained from the *Tetrahymena* Stock Center (tetrahymena.vet.cornell.edu/index). Btu2:mCherry is in the B2086 strain background and Epc1:mCherry is in the SB1969 strain background. Acetylation (*mec17Δ* and *K40R*) and glycylation (*ttll3Δ*) mutant strains were obtained from the *Tetrahymena* Stock Center (Akella et al., 2010, Gaertig et al., 1995, Wloga et al., 2009). Epiplasm mutants (*epc1Δ*) were obtained from (Williams, 2004).

### *Tetrahymena* cell culture

*T. thermophila* strains were cultured in 2% SPP media (2% protease peptone, 0.1% yeast extract, 0.2% glucose, and 0.003% Fe-EDTA) at 30°C unless otherwise indicated. *ttll3Δ*, *epc1Δ*, and corresponding control cells were cultured in MEPP media (2% protease peptone, 2 mM Na_3_ citrate 2H_2_O, 1 mM FeCl_3_, 30 μM CuSO_4_• 5H_2_O, 1.7 μM CaCl_2_; (Orias and Rasmussen, 1976)). Cells collected for analysis were grown to mid-log phase (approximately 3 × 10^5^ cells/mL). Cell counts were determined using a Coulter Counter Z1 (Beckman Coulter). *ttll3Δ* and control (*bld10^mini^Δ*) cells were generated by mixing germline mutant heterokaryon cells of complementary mating types in Dryl’s media for 20 hrs, followed by 7 hrs of recovery in MEPP, and subsequent treatment with Paromomycin (200 μg/mL; (Wloga et al., 2009)). The WT control cells do not have detectable growth, morphology, BB synthesis or BB organization defects.

The forces from ciliary beating were manipulated by increasing temperature and media viscosity (Goto et al., 1982, Galati et al., 2014). For temperature shift experiments, cells were transferred into fresh MEPP media and incubated for 24 hrs at specified temperatures. Ciliary beating was inhibited using 100 µM NiCl_2_ (Bayless et al., 2016, Larsen and Satir, 1991). Media viscosity was increased using 3% polyethylene oxide (PEO, molecular weight 900,000, Acros Organics). PEO was dissolved at 37°C for six hrs and then incubated for 24 hrs at 25°C on a nutator. Cells were suspended in the 3% PEO MEPP media and then cultured at 30°C for 24 hrs (Galati et al., 2014).

### Light microscopy

Immunofluorescence imaging procedures were performed as previously described (Bayless et al., 2016, Thazhath et al., 2002, Williams et al., 1995). For BB visualization, cells were washed in 10mM Tris HCl pH 7.4, followed by PHEM buffer (60 mM 1,4-piperazinediethanesulfonic acid, 25 mM 4-(2-hydroxyethyl)-1-piperazineethanesulfonic acid, 10 mM EGTA, and 2 mM MgCl_2_, pH 6.9), and then fixed in a Triton X-100 / paraformaldehyde solution (0.5% Triton X-100 and 3.2% paraformaldehyde in PHEM buffer) for 5 min. Cells were then washed three times in 0.1% bovine serum albumin (BSA) in PBS (BSA-PBS) before a 24 hr incubation in primary antibody diluted in 1.0% BSA-PBS at 4**°**C. Cells were then washed three times in 0.1% BSA-PBS before a two hr incubation in secondary antibody diluted in 1.0% BSA-PBS at 23°C. Cells were then washed 3 times in 0.1% BSA-PBS and pelleted. 2 µL of cells from the pellet were added to a coverslip and mounted in 6.5 μL Citifluor mounting media (Ted Pella). Samples were then sealed using clear nail polish.

For visualization of microtubules and tubulin PTMs, cells were washed in 10 mM Tris HCl pH 7.4, deciliated using deciliation media (10% Ficoll 400, 10 mM sodium acetate, 10 mM CaCl_2_, 10 mM EDTA, pH 4.2 adjusted with HCl), and incubated for 60 sec. The pH was rebalanced with 15 mM Tris-HCl (pH 7.9), followed by a 10 mM Tris-HCl (pH 7.4) wash, and PHEM (pH 6.9) wash. Cells were permeabilized with 0.5% Triton X-100 in PHEM for 50 sec and then fixed in 2.0% paraformaldehyde. Cells were added to coverslips, dried at 42°C for 20 minutes, rehydrated in 0.1% BSA-PBS for 12 minutes before a 24 hr incubation at 4**°**C with primary antibody diluted in 1.0% BSA-PBS. Cells were then washed three times in 0.1% BSA-PBS before incubation for two hrs at 23°C in secondary antibody diluted in 1.0% BSA-PBS. Cells were washed three times in 0.1% BSA-PBS and mounted with 6.5 µL Citifluor mounting media (Ted Pella). Samples were sealed using clear nail polish. Immunofluorescence for epiplasm was performed as describe previously (Williams et al., 1995). Briefly, cells were incubated on ice for five min, fixed in 35% ethanol + 0.5% TX-100 on ice for 20 min, washed three times in 0.1% BSA-PBS before incubation for one hr in primary antibody at 23°C diluted in 1.0% BSA-PBS, and then washed three times in 0.1% BSA-PBS before incubation for one hr in secondary antibody at 23°C diluted in 1.0% BSA-PBS. Cells were then washed three times in 0.1% BSA-PBS before mounting with 6.5 µL Citifluor mounting media (Ted Pella). Samples were sealed using clear nail polish.

The primary antibodies used in this study were α-TtCen1 (1:1000; (Stemm-Wolf et al., 2005), α-monoglycylation (1:500, TAP952, EMD Millipore; (Callen et al., 1994)), α-polyglycylation (1:500, AXO49, EMD Millipore; (Callen et al., 1994)), α-glutamylation (1:1000; GT335; Adipogen; (Wolff et al., 1992)), α-striated fiber (1:1000; 5D8; (Jerka-Dziadosz et al., 1995)), α-α-tubulin (1:200; 12G10; AB_1157911), α-acetylated tubulin (1:200; 6-11 B-1; (LeDizet and Piperno, 1991)), α-Epc1 (1:500; 8G1; (Williams et al., 1995)). Secondary antibodies (Alexa Fluor 488-, 594-, or 647-goat α–rabbit IgG, Alexa Fluor 488-, 594-, or 647-goat α–mouse IgG (Invitrogen)) were used at 1:1000.

Fluorescence imaging was performed as previously described (Bayless et al., 2016). Briefly, either a Nikon wide-field TiE fluorescence microscope stand equipped with a 100× NA 1.4 Plan Apo objective and an Andor Xyla 4.2 CMOS camera or a Nikon swept-field confocal TiE fluorescence microscope stand equipped with a 100× NA 1.4 Plan Apo objective and an Andor iXon EMCCD camera were used. Images were acquired using NIS Elements software. All fluorescence imaging was conducted at 25°C and exposure times were between 50-500 msec, depending on the experiment. For experiments requiring quantitative comparisons of fluorescence intensity, matched samples were imaged on the same day with the same acquisition settings and representative images were scaled equally.

### Quantification of BB parameters

#### BB-appendage microtubule elongation and length

Images acquired for analysis were from samples with matched experimental, preparation, and image acquisition conditions. BB-appendage microtubule length and elongation was measured as the distance from the approximated centroid of Btu2-mCherry BB signal to the tips of BB-appendage microtubules minus the half-width half-maximum of the BB signal (HWHM). BB-appendage microtubule extension from the BB in WT and *ttll3Δ* cells was measured as the distance from BB centroid (brightest Centrin pixel after application of gaussian blur) to the approximated tip of the microtubule bundle (12G10, α-tubulin). Measurements were performed using Image J (NIH image).

#### Total cell fluorescence

Full cell Z-stacks were sum projected and were background subtracted using ImageJ. The corrected total cell fluorescence intensity was obtained by measuring the integrated intensity of the total outlined cell and background subtracted using the integrated intensity of a region outside of the cell.

#### BB disorganization

Disorganized BBs were identified as any BB with an approximated centroid deviating by greater than 1.0 μm from the boundary of the BB row. The BB row boundary was defined by the edges of the nearest organized anterior and posterior BBs. BBs that were 1.0 μm outside the bounds of a row were considered disorganized. Only BBs from the medial and anterior portions of the cell were counted due to the difficulty of assigning BB rows in the posterior end of the cell where BBs are less dense and have less defined positioning.

#### BB attachment to cortex

BBs were considered cortically detached if they were greater than 1.0 μm interior to the adjacent cortically attached BBs. BB positioning was also measured relative to the cell cortex using epiplasm (Epc1:mCherry) and DIC imaging.

### Electron Tomography

Cells were prepared for electron tomography as previously described (Giddings et al., 2010, Meehl et al., 2009). Cells were gently spun into 15% dextran (molecular weight 9000–11,000; Sigma-Aldrich) with 5% BSA in 2% SPP. A small volume of concentrated cells was transferred to a sample holder and high-pressure frozen using a Wohlwend Compact 02 high pressure freezer (Technotrade International). After low-temperature freeze substitution in 0.25% glutaraldehyde and 0.1% uranyl acetate in acetone, cells were slowly infiltrated with Lowicryl HM20 resin. Serial thick (250–300 nm) sections were cut using a Leica UCT ultramicrotome. The serial sections were collected on Formvar-coated copper slot grids and poststained with 2% aqueous uranyl acetate for four min followed by Reynold’s lead citrate for three min.

Dual-axis tilt series (−60 to +60°) of *Tetrahymena* cells were collected on a Tecnai F30 intermediate voltage electron microscope (FEI). Images were acquired using the SerialEM acquisition program with a Gatan CCD camera at 1.2 nm / pixel. Serial section tomograms of *Tetrahymena* cortical structures were generated using the IMOD software package (Giddings et al., 2010, Kremer et al., 1996, Mastronarde, 1997). In total, eight tomograms were reconstructed. Three-dimensional (3D) models were created using the IMOD software package (bio3d.colorado.edu/imod/). Diagrams of 3D structures in Fig. 1D were generated using Blender Software (blender.org).

### Statistical Analyses

All experimental data sets represent a minimum of three biological replicates. The total number of cells and structures analyzed for each data set is described in the Figure Legends. Statistical tests were run in Prism8 (GraphPad Software). Categorical data sets were analyzed using the Fischer’s exact test. Normally distributed continuous data sets were analyzed using the Student’s t-test. Non-normally distributed continuous data sets were analyzed using the Mann-Whitney test and Kruskal-Wallis test (multiple comparisons). Unless otherwise noted, all p values are numerically presented. Circular and directional statistical tests and plots were generated using ORIANA software (Kovach) to calculate R-values (mean-vector). All error bars in bar graphs and x-y plots indicate standard error of the mean (SEM). Lines and bars on all dot plots indicate the mean and standard deviation, respectively.

## ACKNOWLEDGEMENTS

The authors would like to thank Alexander Stemm-Wolf and Marisa Ruehle for critical reading and feedback for this manuscript. We appreciate the discussions and expertise provided by the Pearson Lab. The authors would also like to thank the *Tetrahymena* Stock Center (Cornell University) for strains and information for culturing *Tetrahymena*. Specimen preparation and electron tomography was performed by Garry Morgan and Eileen O’Toole in the Boulder Electron Microscopy Services at the University of Colorado, Boulder.

## COMPETING INTERESTS

No competing interests declared.

## FUNDING

This work was supported by the National Institutes of Health (NIH-NIGMS RO1GM127571 to MW, NIH-NICHD R21HD092809 to JG, and NIH-NIGMS R01GM099820 to CGP) and the American Cancer Society (ACS-RSG-16-157-01-CCG to CGP).

